# Prevalence of pulmonary tuberculosis and associated factors among prisoners in Western Oromia, Ethiopia: A cross-sectional study

**DOI:** 10.1101/869727

**Authors:** Keneni Ephrem Dibissa, Zelalem Desalegn Waktole, Belachew Etana Tolessa

## Abstract

**Background:** Prisoners are a disproportionately at high risk for tuberculosis. This is because; prisons represent dynamic communities where at-risk groups congregate. It increases the transmission rate because of overcrowding and living together with infected individuals. This study was done to determine the prevalence of pulmonary tuberculosis and associated factors among prisoners of Western Oromia, Ethiopia in 2017.

**Methods:** A cross-sectional study was conducted among prisoners who have a history of cough for two weeks or more. Data were collected from 270 participants and sputum sample was collected from 249 prisoners and analyzed in GeneXpert for having pulmonary tuberculosis. Logistic regression analysis was used to identify factors associated with the development of pulmonary tuberculosis among prisoners.

**Results:** The overall prevalence among suspected cases was (15.6%; 95% CI (11.5, 20)) which makes the point prevalence of pulmonary tuberculosis were 744 per 100,000 of prisoners. Prisoners who had history of cigarette smoking before imprisonment (AOR=3.55; 95% CI (1.29, 9.78)), contacted with known TB patient (AOR=5.63; 95% CI (2.19, 14.41)), share prison cell with TB patients (AOR=3.51; 95% CI (1.34, 9.19)) and Body Mass Index <18.5kg/m2 (AOR=8.87; 95% CI (3.23, 24.37)) were more likely to have pulmonary tuberculosis.

**Conclusion:** A higher prevalence of pulmonary tuberculosis was observed among prisoners in the three prisons of Wollega Zones. To avert this problem, screening of prisoners should be done at the entry and separation of inmates with symptoms of tuberculosis should be done.

## Introduction

Tuberculosis (TB) is an airborne infectious disease caused by bacillus mycobacterium tuberculosis. It typically affects the lungs (pulmonary Tuberculosis), but it can also affect other sites (extra pulmonary Tuberculosis). The disease spreads in the air when sick people expel the bacteria while talking, coughing, singing, sneezing and spitting [1].

About 95% of all cases and 99 % of deaths occur in low and middle-income countries. Twenty-two countries contribute to 99% of the world’s TB burden. Besides this, household costs of TB are substantial estimates suggest that tuberculosis costs the average patient three or three months of lost earnings, which can represent up to 30 percent of annual household income [2, 3].

TB mostly affects socially marginalized and other poor high-risk groups such as prisoners, intravenous drug users, migrants, and poor socio-economic groups. Prison inmates constitute a high risk-group for tuberculosis (TB) in both developing and industrialized countries [4]. In terms of cases, the best estimates for 2015 are that there were 10.4 million new TB cases (including 1.2 million among HIV-positive people), of which 5.9 million were among men, 3.5 million among women and 1.0 million among children. Overall, 90% of cases were adults and 10% of children [5].

Tuberculosis (TB) has remained a major global health problem. In 2015, it was one of the top 10 causes of death worldwide, ranking above HIV/AIDS an infectious disease. Without treatment, about 70% of people with sputum smear-positive pulmonary TB will die within 10 years after infection. Ethiopia is one of the twenty-two countries affected by a high burden of TB. According to the World Health Organization report (WHO), these countries have about 200,000 new TB cases. From these, Ethiopia ranked tenth among the world’s 22 high burden countries [5, 6].

Poverty, malnutrition and over-crowded living conditions have been known for decades to increase the risk of developing the disease. Prisons are often high-risk environments for TB transmission because of severe overcrowding, poor nutrition, poor ventilation, and limited access to often health care. Moreover, prisoners are overwhelmingly male, are typically aged 15-45 years which study indicated high risk for TB [6]. Furthermore, Most of the prisoners are predominantly from poorly educated and socioeconomically deprived sectors of the population where TB infection and transmission are higher [6].

In addition to overcrowding, long prison stays, low Body Mass Index (BMI), previous TB treatment, loss of appetite, poor nutrition, and HIV infection have been documented as risk factors for TB [7, 8]. Because of these, prison is also specifically taken as a risky place for the transmission of TB in low and middle-income countries. Its prevalence is estimated to be about ten to a hundredfold in the prison than the general population [9, 10].

Studies that indicated the prevalence of TB in the prison have reported higher prevalence in Ethiopia. The study was done in Southern Ethiopia, Gamo Gofa prison, which indicated about 3.2% prevalence of TB among inmates. Another study was done in the Hadiya Zone; south Ethiopia also showed a three times higher prevalence of TB than in the general population [11, 12]. But there is limited evidence regarding the TB prevalence in Western Ethiopia.

This study generated information on the current prevalence of TB among prisoners in Wollega Zones, western Ethiopia. The study was also aimed to identify factors associated with TB infection in prisons. This will help TB programmers and authorities in re-designing the existing programs for TB prevention in prisons. Thus, there are needs to accurately define specific factors driving TB and the prevalence of the problem among prisoners of Wollega Zones.

## Methods and materials

A cross sectional study design was conducted in three prisons found in Western Oromia Region, Ethiopia. These prisons are located in Gimbi, West Wollega; Nekemte, East Wollega; and Dambi Dollo, Kelem Wollega. The prisons have a total population of 2, 228, 1969 and 1, 447 respectively during study period. 270 prisoners were presented with cough and other suggestive symptoms during screening and all of them were included in the study. Based on this, 108 prisoners from Gimbi, 97 prisoners from Nekemte and 71 prisoners from Dambi Dollo were included.

Participants of the study were selected based on their history of cough of two weeks and above before data collection date. Then, all prisoners with history of cough were screened for the presence of TB using GeneXpert. Beside this, PTB patients who were taking anti-TB treatment were included in the study. HIV positive inmates with cough of any duration were also included in this study. However, prisoners who were unable to produce sputum and unable to communicate during data collection were excluded. Then, all prisoners with cough of two weeks or more were screened for having pulmonary TB through mass screening strategy.

Data were collected by using interviewer administered questionnaire that was initially adapted from previous studies [14]. The questionnaire was prepared in English and translated into ‘Afaan Oromo’ (regional working language) for data collection. It has sections on socio-demographics, imprisonment history and morbidity history of prisoners. Data was collected by six nurses and three laboratory technologists. One nurse and one laboratory technologist supervised the overall data collection process.

Before data collection, prisons’ health committees and health professionals working in prisons were oriented on the purpose of the research. Then, all inmates were registered with support of the health committee members in each prison.

### Sputum Collection and transportation

A single sputum specimen is recommended for GeneXpert. The sputum was collected outside the laboratory or clinics in a well-ventilated space. The prisoners were instructed to produce sputum through coughing to get sputum from lower respiratory organs. Respondents were also asked to wash their mouth using clean water before the sputum was taken.

Then, they were asked to inhale deeply 2–3 times and breathe out strongly each time cough deeply from chest to produce sputum. Sputum specimens were collected in “Falcon tube” with 30-50 ml capacity. This was because it was translucent and has walls that allow easy labeling. Then, the sputum was transported using triple package to hospital for laboratory to be evaluated. Appropriate procedure of using the GeneXpert test was followed during laboratory test [15].

### Weight and height measurements

Body weight was determined to the nearest 0.1kg on weight scale and height was measured to the nearest 0.1cm. Then body mass index (BMI) was calculated as weight in kilogram of the individual divided by the square of the height in meter. Then nutritional status of determined as severe malnutrition (BMI <15.9 kg/m2), moderate malnutrition (BMI= 16–16.9 kg/m2), mild malnutrition (BMI =17–18.4 kg/m2) and well-nourished for BMI >18.4 kg/m^2^ [16].

### Data quality control

Data collectors and supervisors were trained for one day. Then, data collection tools were pre-tested on 5% of the sample on Shambu town prison. The GeneXpert Diagnostic System automatically performs internal quality control for each sample. GeneXpert has an in built internal quality control system within the test cartridge. In addition, laboratory workers had checked new batch of cartridges with known positive and negative specimens before using them for patient sample testing.

### Data analysis

The data were entered and analyzed using SPSS version 20. Statistical summary measures like frequency, percentage and mean with standard deviation were calculated to describe the data. Then, bivariate logistic regression analysis was employed to check crude association between pulmonary tuberculosis status and explanatory variables. Based on this, variables with p-value <0.25 were entered to multivariable logistic regression to identify the factors that affect pulmonary tuberculosis. Odds ratio and corresponding 95% confidence intervals were used to quantify the degrees of association between independent variables and PTB status. All associations with p-value ≤0.05 were considered as being statistically significant.

### Operational definitions

**TB Positive screen:** identifying a person with symptoms and findings consistent with tuberculosis involves screening of patients with particular attention to cough of two weeks or more duration.

**Suggestive symptoms:** that help to identify presumptive TB include fever, night sweating, and weight loss.

**New cases of TB**: those patients who have never been diagnosed for TB before but diagnosed by GeneXpert during data collection.

**Existing TB case:** refers to a patient who had been diagnosed for TB and receiving anti-TB drugs during data collection.

**Malnutrition:** a person with Body Mass Index (BMI) less than18.5kg/m2.

**Good housing ventilation condition:** if the house has adequate windows and doors to ventilate the rooms with fresh air observed by data collector during data collection period

**Good housing ventilation condition:** if the house has no adequate windows and doors to ventilate the rooms with fresh air observed by data collector during data collection period

**Lower respiratory organs:** respiratory organs including trachea, bronchi, bronchial, alveoli and the whole lunge

### Ethical statement

Ethical clearance letter was obtained from ethical review committee of Wollega University and was brought to the administrative bodies of prisons, Zonal health department, Gimbi General Hospital, Nekemte Referral Hospital and Dambi Dollo Hospital to get permission for the study and material support. Before data collection written consent was taken from study participants. Consent was obtained from a parent or guardian on behalf of any participants under the age of 18. Confidentiality was insured and maintained by the investigators and data collectors.

## Results

### Characteristics of study participants

A total of 5, 644 prisoners 5441 (96.4%) males and 203 (3.6%) females were screened and 644 had history of cough. Among those who were coughing 249 were with cough greater than or equal to 2 weeks and 395 were with cough less than 2 weeks. In this study 249 with cough greater than or equal to 2 weeks and the existing 31 PTB patients on treatment were participated. From these, 270 prisoners majority of them were male (97.4%), married (57.4%) and attended primary school (53.7%). Nearly half (42.6%) of the suspected PTB case-patients had history of smoking with a median duration of smoking of 7 years before imprisonment and 111(41.1%) have history of chat chewing (Table1).

**Table 1:** Characteristics of the study population among prisoners in Wollega zones, Western Oromia, Ethiopia, 2017 (n=270)

| Variables | Frequency | Percent |
| --- | --- | --- |
| **Sex** |  |  |
| Male | 263 | 97.4 |
| Female | 7 | 2.6 |
| **Age** |  |  |
| 15-24 | 102 | 37.8 |
| 25-34 | 84 | 31.1 |
| 35-44 | 39 | 14.4 |
| >45 | 45 | 16.7 |
| **Religion** |  |  |
| Muslim | 49 | 18.1 |
| Protestant | 134 | 49.6 |
| Orthodox | 72 | 26.7 |
| Adventist | 14 | 5.2 |
| Other | 1 | 0.4 |
| **Ethnicity** |  |  |
| Oromo | 246 | 91.1 |
| Amhara | 18 | 6.7 |
| Others | 6 | 2.3 |
| **Marital status** |  |  |
| Single | 100 | 37 |
| Married | 155 | 57.4 |
| Others | 15 | 5.5 |
| **Educational status** |  |  |
| No formal education | 81 | 30 |
| Primary school (1-8) | 145 | 53.7 |
| Secondary school (9-12) | 35 | 13 |
| College and /or university (12+) | 9 | 3.3 |
| **Occupation before imprisonment** |  |  |
| Government employee | 9 | 3.3 |
| Farmer | 152 | 56.3 |
| Merchant | 25 | 9.3 |
| Daily labor | 29 | 10.7 |
| Student | 43 | 15.9 |
| Other | 13 | 2.4 |
| **Residence** |  |  |
| Rural | 203 | 75.2 |
| Urban | 67 | 24.8 |
| **Cigarette smoking** |  |  |
| Yes | 115 | 42.6 |
| No | 155 | 57.4 |
| **‘Khat’chewing** |  |  |
| Yes | 111 | 41.1 |
| No | 159 | 58.9 |

Regarding prison related history of participants, 179 (66.3%) had stayed for less than two years in current prison, 93 (34.4%) were sharing cell with TB patients. About 160 (59.3%) of the study participants had stayed for greater or equal to three weeks with person who have persistent cough. Furthermore, about 154(57%) of the study participants share drinking and eating materials with other prisoners. There were 153(56.7%) prisoners who had no support from family in terms of visit and bringing food and 218(80.7%) lives in a prison which has greater than 100 prisoners per cell (Table 2).

**Table 2:** Prison related characteristics of the study population in prisons of Wollega zones, Western Oromia, Ethiopia, 2017 (n=270)

| Character/variables | Labels | Frequency | Percent |
| --- | --- | --- | --- |
| Duration of stay in current prison | ≤24 months | 179 | 66.3 |
|  | >24 months | 91 | 33.7 |
| Sharing cell with TB Patient | Yes | 93 | 34.4 |
|  | No | 177 | 65.6 |
| Housing ventilation condition | Good | 89 | 33 |
|  | Bad | 181 | 67 |
| Duration of stay with Person who have persistent cough | <3 weeks | 107 | 39.6 |
|  | ≥3 weeks | 160 | 59.3 |
|  | I don't know | 3 | 1.1 |
| Sharing drinking and eating materials with other persons | Yes | 154 | 57 |
|  | No | 116 | 43 |
| Support from family in terms of visit and bringing food | I don't have | 153 | 56.7 |
|  | visit only | 18 | 6.7 |
|  | Food only | 3 | 1.1 |
|  | visit and food | 96 | 35.6 |
| Prisoners per cell | ≤100 | 52 | 19.3 |
|  | >100 | 218 | 80.7 |

From total study participants 133(49.3%) had received treatment for their current complaint and 32 (11.9%) from prison’s clinic, 16(5.9%) other health institutions and 85(31.5%) from both. About 70(25.9%) study participants had contact with known TB patient at home and 56(20.7%) had history of hospitalization. 27(10%) of them were diagnosed for Diabetic mellitus/Hypertension and 19(7%) received treatment for different chronic diseases. 60(22.2%) of those study participants had Body Mass Index (BMI) < 18.5 kg/m2 and 3(1.1%) were HIV positive (table 3).

**Table 3.** Morbidity related characteristics of study participants in Wollega zones prison western Oromia, Ethiopia, 2017.

| Character/variables | Labels | Frequency | Percent |
| --- | --- | --- | --- |
| Treatment for your current complaint | Yes | 133 | 49.3 |
|  | No | 137 | 50.7 |
| Place of treatment for current symptom | Health institution | 16 | 5.9 |
|  | Prison's clinic | 32 | 11.9 |
|  | Both | 85 | 31.5 |
| History of contact with known TB patient at home | Yes | 70 | 25.9 |
|  | No | 200 | 74.1 |
| History of hospitalized | Yes | 56 | 20.7 |
|  | No | 214 | 79.3 |
| Diagnosed for Diabetic mellitus/Hypertension | yes | 27 | 10 |
|  | No | 225 | 83.3 |
|  | I don't know | 15 | 5.6 |
| Treatment for chronic diseases | Yes | 19 | 7 |
|  | No | 67 | 24.8 |
| Body mass index | < 18.5 kg/m2 | 60 | 22.2 |
|  | ≥ 18.5kg/m2 | 210 | 77.8 |
| HIV test result | Positive | 3 | 1.1 |
|  | Negative | 267 | 98.9 |

### Prevalence of pulmonary TB among prisoners

Out of those 249 prisoners who provided sputum, 11(4.1%) of them were found with new cases of TB. In addition to the previous cases being treated, the overall prevalence of pulmonary tuberculosis (TB) among suspected cases was 15.6 % with 95% CI (11.5-20) in prisons of Wollega zones. This will result in 744 cases of pulmonary TB per 100,000 prisoners. Majority of cases were found in Gimbi prison (14 existing and 6 new cases). Among the study participants Mycobacterium tuberculosis (MTB) were detected in (8.9%) with two weeks of cough, (19.7%) with cough of four weeks and (21.4%) with cough of six weeks (Table 4).

**Table 4:**
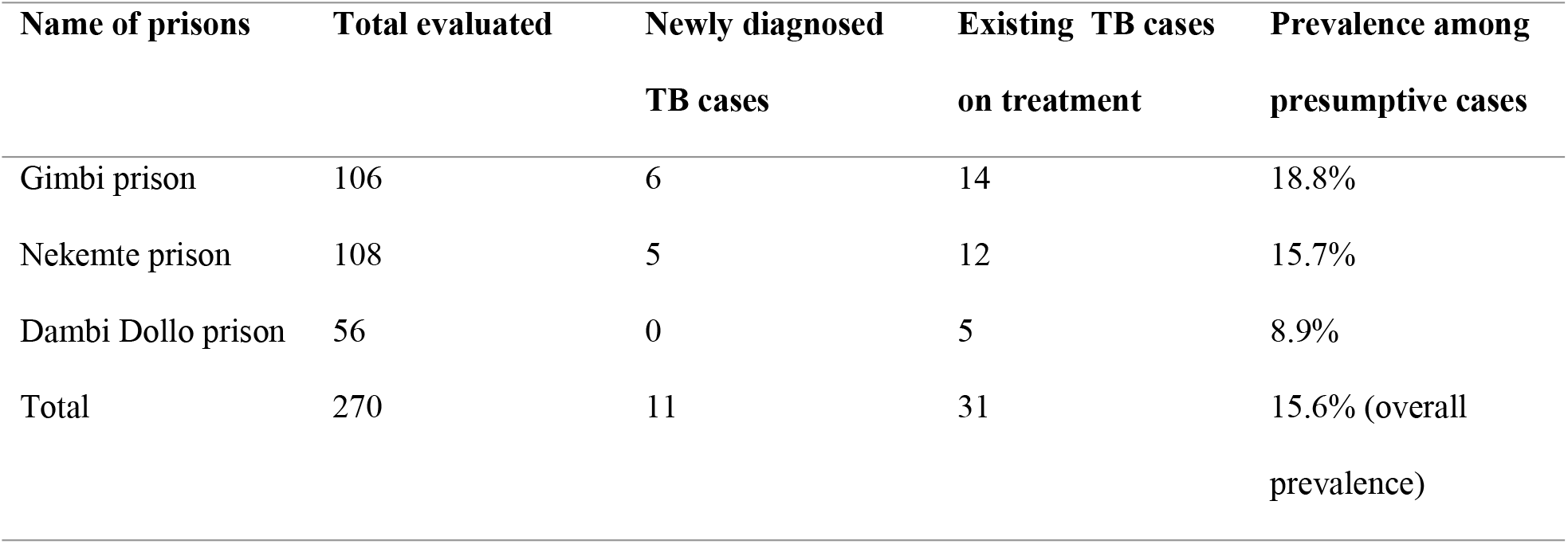
Prevalence of TB among the study population in prisons of Wollega zones, Western Oromia, Ethiopia, 2017 (n=270)

### Factors associated with pulmonary TB

The study found that, prisoners who smoke cigarette before imprisonment (AOR=3.56; 95% CI (1.29, 9.78)), history of contact with known TB patient at home (AOR=5.63; 95% CI (2.19, 14.41)), longer duration of stay in current prison (AOR=3.21; 95% CI (1.12, 9.17)) and low Body Mass Index <18.5kg/m^2^ (AOR= 8.87; 95% CI (3.23, 24.37)) were associated with prevalence of TB (Table 5).

**Table 5:**
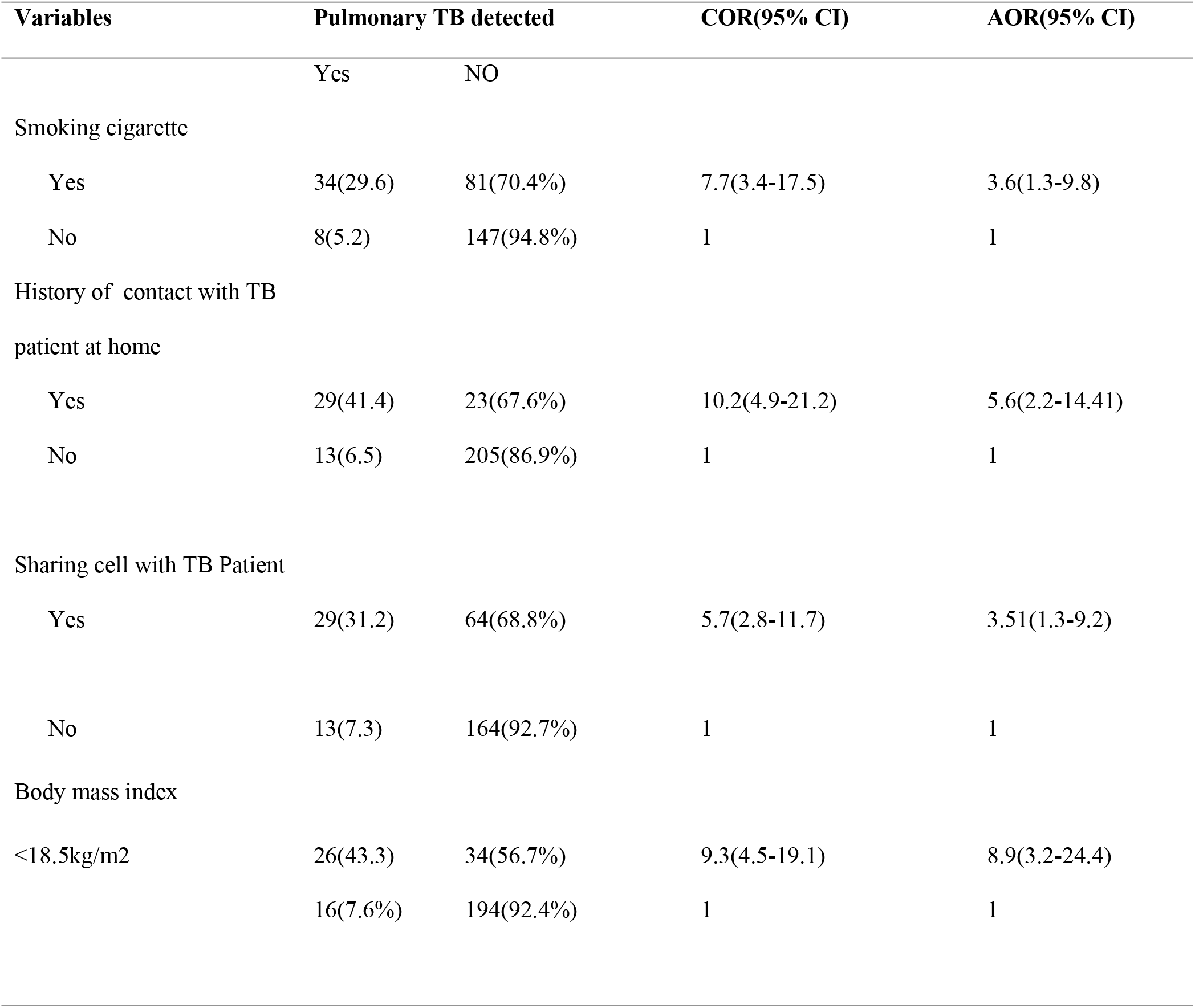
Multivariate analyses of factors associated with PTB among prisoners of Wollega Zones, Western Oromia, Ethiopia, 2017.

## Discussion

This study showed that the prevalence of Pulmonary TB in prison is about 744 cases per 100,000 prisoners. Furthermore, the magnitude of TB among suspects was 15.6 % (95% CI (11.5, 20)). This finding indicated higher prevalence when compared to the southern Ethiopia, Gamo Gofa which has prevalence 623 per 100,000 prisoners. It is also higher than the study done in Eastern Ethiopia (9%) and North Gondar prison (10.4%).

The observed prevalence is 7 times higher than in general population. New cases of TB were also observed during data collection which indicated undiagnosed TB in the prisons which are source of infection for other prisoners. Because of this, there is a need for regular screening of TB during intake on routine basis. The use of GeneXpert may also contribute for better case detection due to that GeneXpert is more accurate and reliable than sputum smear microscopy in predicting pulmonary TB [17].

Smoking was significant risk factor of PTB in the current study. This finding is similar with the study conducted in Tanzania [18], and southern Ethiopia [14]. On bivariate analysis participants who had history of chewed ‘chat’ before imprisonment had association and the risk of developing PTB positive among ‘chat’ chewers were 5 times than those who were not chewed ‘chat’ which is not reported in any other previous study done in Ethiopia like, Hadiya, Wolaita and Bedelle [12, 13, 14], which could be due to high numbers of chewers in current study areas but on multivariate analysis it had no association with PTB with (AOR=1.33, 95% CI (0.43, 4.15)).

Sharing prison cell also significantly associated with TB infection which is in line with the study conducted in Eastern Ethiopia [11]. This is because when the non-infected individuals share room there is easy inhalation of droplets from infected individuals. Having history of contact with known tuberculosis patient before imprisonment had significantly associated with acquiring pulmonary tuberculosis in the current study (AOR=5.63, 95% CI (2.19, 14.41)); this significance was seen also in study conducted in Wolaita [14]. This association could be due to reactivation of latent tuberculosis infection because of immunity degradation and person to person transmission is also more likely with proximity with the infected person.

In the current study, nutritional status (BMI) was significantly associated with pulmonary tuberculosis positivity (AOR=8.87, 95% CI (3.23, 24.37)). BMI <18.5kg/m^2^ had causative effect on PTB as compared to those with BMI ≥ 18.5kg/m^2^ which is in agreement with study conducted in Bedelle [13], mentioned low BMI (i.e. ≤ 18.5kg/m^2^) as the risk factor for TB. The possible reason for this could be malnutrition is equally recognized as a risk factor for the reactivation of latent TB and its progression to disease. This assumption is supported by previous studies in Hadiya [12], this link is besides bi-directional as TB can cause or predispose to malnutrition. Hence, when comparing individuals with and without active PTB in this cross-sectional study with respect to their nutritional status, a causal effect cannot be assigned to malnutrition, even though it came out as a significant predictor in the multivariate analysis.

## Conclusions

We can conclude from this study that the prevalence of PTB in prisons of Wollega Zones’ is high and there were high burden of undetected and infectious PTB cases in the prisons. History of cigarette smoking before imprisonment, history of contact with known TB patients at home and body mass index were identified predictors of PTB among prisoners in Wollega Zones. In brief, the high prevalence and associated risk factors for PTB may favor an active transmission of PTB and put the prison population at increased risk of developing PTB. This could be also a great health threat to the surrounding community.

Based on the above results, we would like to recommend the following to all concerned bodies. Conducting an entry and exit screening of prisoners to identify early infectious cases, prevent further delay in diagnosis and reduce prolonged transmission of TB in the prisons. Provide adequate and diversified food for prisoners and adequate ventilation. Segregation of smear positive patients for the full initial phase of directly observed treatment (DOTS) treatment should be given a priority in prisons’ TB control strategies and continuous health information on mode of PTB and its prevention mechanisms. Strengthen health committee to identify and screen PTB positive among suspected cases. Periodic mass screening and follow up of those PTB patients by knowing their addresses when they released from prison in order to know their treatment outcome, undergo supportive supervision of prisons’ health workers and monitoring of TB prevention and care services in prisons.

## Availability of data and materials

All the data and materials are available with the authors.

## Authors’ contributions

KE has been involved in conception, writing the study protocol, formulating the study design, and training of data collectors, data entry, analysis and interpretation of data. ZD participated in design, interpretation of data, reviewing intellectual content; supervise overall process of the process and manuscript preparation. BE was part of data quality check, providing important comments, supervise overall process and review manuscript.

## Acknowledgements

We would like to thank Wollega University, Gimbi General Hospital, Nekemte Referral Hospital, Dambi Dollo Hospital and West Wollega Zone Health Office for material support. We are also grateful to the data collectors, supervisors and study participants for their voluntarily participation.

